# Implications of gene tree heterogeneity on downstream phylogenetic analyses: A case study employing the Fair Proportion index

**DOI:** 10.1101/2023.01.21.525012

**Authors:** Kristina Wicke, Md Rejuan Haque, Laura Kubatko

## Abstract

Many questions in evolutionary biology require the specification of a phylogeny for downstream phylogenetic analyses. However, with the increasingly widespread availability of genomic data, phylogenetic studies are often confronted with conflicting signal in the form of genomic heterogeneity and incongruence between gene trees and the species tree. This raises the question of determining what data and phylogeny should be used in downstream analyses, and to what extent the choice of phylogeny (e.g., gene trees versus species trees) impacts the analyses and their outcomes.

In this paper, we study this question in the realm of phylogenetic diversity indices, which provide ways to prioritize species for conservation based on their relative evolutionary isolation on a phylogeny, and are thus one example of downstream phylogenetic analyses. We use the Fair Proportion (FP) index, also known as the evolutionary distinctiveness score, and explore the variability in species rankings based on gene trees as compared to the species tree for several empirical data sets. Our results indicate that prioritization rankings among species vary greatly depending on the underlying phylogeny, suggesting that the choice of phylogeny is a major influence in assessing phylogenetic diversity in a conservation setting.

While we use phylogenetic diversity conservation as an example, we suspect that other types of downstream phylogenetic analyses such as ancestral state reconstruction are similarly affected by genomic heterogeneity and incongruence. Our aim is thus to raise awareness of this issue and inspire new research on which evolutionary information (species trees, gene trees, or a combination of both) should form the basis for analyses in these settings.

## Introduction

Estimating the evolutionary relationships among organisms is a central goal in evolutionary biology. On one hand, phylogenies are inferred to uncover past evolutionary trajectories and understand the relatedness and differences among organisms. On the other hand, phylogenies provide the basis for downstream phylogenetic analyses such as studying trait evolution (see, e.g., the review by [16]), ancestral state reconstruction (see, e.g., [8] for an overview and [18] for a more recent study), estimation of diversification rates and testing of macroevolutionary models (e.g., [14, 19, 10]), and quantifying biodiversity (e.g., [3, 6, 31]).

To provide meaningful results, these downstream analyses rely on accurate phylogenies representing the evolutionary relationships among the organisms studied. Traditionally, phylogenetic trees have been inferred from single genes and the resulting gene trees were assumed to be a valid estimate for the species tree, i.e., the “true” evolutionary history of the species under consideration. However, the advent of whole genome sequencing has resulted in increased appreciation of the differences between these gene trees and the species tree. Discordance among gene trees and between gene trees and the species tree is known to arise from numerous biological processes, including incomplete lineage sorting, lateral gene transfer, and gene duplication and loss (e.g., [11, 15, 17]). Downstream phylogenetic analyses thus face the challenge of conflicting signal in the data and the question of which evolutionary information (species trees, gene trees, or a combination of both) to employ in these settings.

In this paper, we exemplarily study the effects of phylogenetic variation in the form of gene tree heterogeneity in the context of phylogenetic diversity conservation. Phylogenetic diversity (PD) is a quantitative measure of biodiversity introduced by [3]. Given a weighted phylogeny, it sums the branch lengths connecting a subset of the species, thereby linking evolutionary history to feature diversity [3]. In fact, preserving PD and the “Tree of Life” has become an integral part of conservation considerations (see, e.g., the “Phylogenetic Diversity Task Force” initiated by the IUCN [7]).

While PD captures the biodiversity of sets of species, evolutionary isolation metrics, also known as phylogenetic diversity indices, apportion the total diversity of a tree among its leaves and quantify the relative importance of species for overall biodiversity based on their placement in the tree. Various methods can be devised to distribute the total diversity of a tree across present-day species, and a variety of phylogenetic diversity indices for phylogenetic trees has been introduced in the literature (for an overview, see, e.g., [31, 23]). One of the most popular indices is the *Fair Proportion* (FP) index (also known as *evolutionary distinctiveness* (ED) score) introduced by [22, 6]. Its underlying idea is to assign each species a “fair proportion” of the total PD of a tree. More precisely, each species descended from a given branch in the phylogeny receives an equal proportion of that branch’s length.

The FP index has been employed in the EDGE of Existence programme, a global conservation initiative focusing specifically on threatened species that represent a significant amount of unique evolutionary history ([6]; see also https://www.edgeofexistence.org/). This project aims at identifying species that are both **e**volutionary **d**istinct and **g**lobally **e**ndangered, where evolutionary distinctiveness is measured in terms of the FP index and the globally endangered score is based on the IUCN Red List Category. However, because the FP index heavily depends on the underlying phylogenetic tree and its branch lengths, different phylogenies for the same set of species may result in different prioritization orders. In this note, we show that this is indeed a widespread phenomenon occurring across various groups of organisms by analyzing nine multilocus data sets from the literature and comparing the FP prioritization orders obtained on individual gene trees, on species trees, and by averaging over gene trees.

We remark that while the FP index is currently employed in the EDGE of Existence project, a revised “EDGE2 protocol” has recently been advocated for in the literature in order to overcome some of the short-comings of the existing approach [5]. The new protocol is more comprehensive and for instance incorporates uncertainty in the phylogeny and extinction risks of species as well as PD complementarity [4]. In particular, it replaces the FP index by an “ED2” score which is equivalent to the so-called “heightened evolutionary distinctiveness (HED)” score [29]. Intuitively speaking, the ED2 score of a species takes into account both the species’ placement in an underlying phylogeney as well as the extinction risk of its close relatives. Importantly, however, the ED2 score also relies on a given phylogeny as a precursor and is thus likely also affected by genetic heterogeneity and incongruence between gene trees and the species tree. In this study, we thus employ the simpler FP index to illustrate the effects of phylogenetic variation on species prioritization. However, we emphasize that we employ the FP index primarily as an example of a phylogenetic downstream analysis. While we analyze nine multilocus data sets from the literature covering a wide range of organisms, our study does not imply that these species currently deserve or do not deserve conservation attention. Moreover, while EDGE-like studies typically use dated and ultrametric phylogenies [5], we do not have enough information (e.g., about the fossil record) for the nine multilocus data sets we analyze to date them (however, as described below, we enforce a molecular clock for all species trees but not gene trees). Additionally, phylogenetic conservation studies are ideally performed using large phylogenies with near-complete taxonomic groups, which our data sets unfortunately also lack. Our case study is thus meant as an illustration of the impacts of phylogenetic heterogeneity on downstream analyses in general and not as a real-world conservation study.

## Materials and Methods

### The FP index

Let *T* be a rooted phylogenetic tree with leaf set *X* = {*x*_1_, …, *x*_*n*_} and root *ρ*, where each edge *e* is assigned a non-negative length *l*(*e*) *∈* ℝ_*≥*0_. Then, the FP index [22, 6] for leaf *x*_*i*_ *∈ X* is defined as 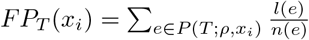, where *P* (*T* ; *ρ, x*_*i*_) denotes the path in *T* from the root *ρ* to leaf *x*_*i*_ and *n*(*e*) is the number of leaves descended from *e*. As an example, for tree *T* depicted in Fig. 1 and leaf *x*_1_, we have 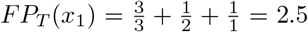. Analogously, *FP*_*T*_ (*x*_2_) = 3.5, *FP*_*T*_ (*x*_3_) = 2, *FP*_*T*_ (*x*_4_) = 3, and *FP*_*T*_ (*x*_5_) = 3. Note that the FP index induces a natural ordering of species, where species with the larger values of the FP index are given higher conservation priority.

**Figure 1:**
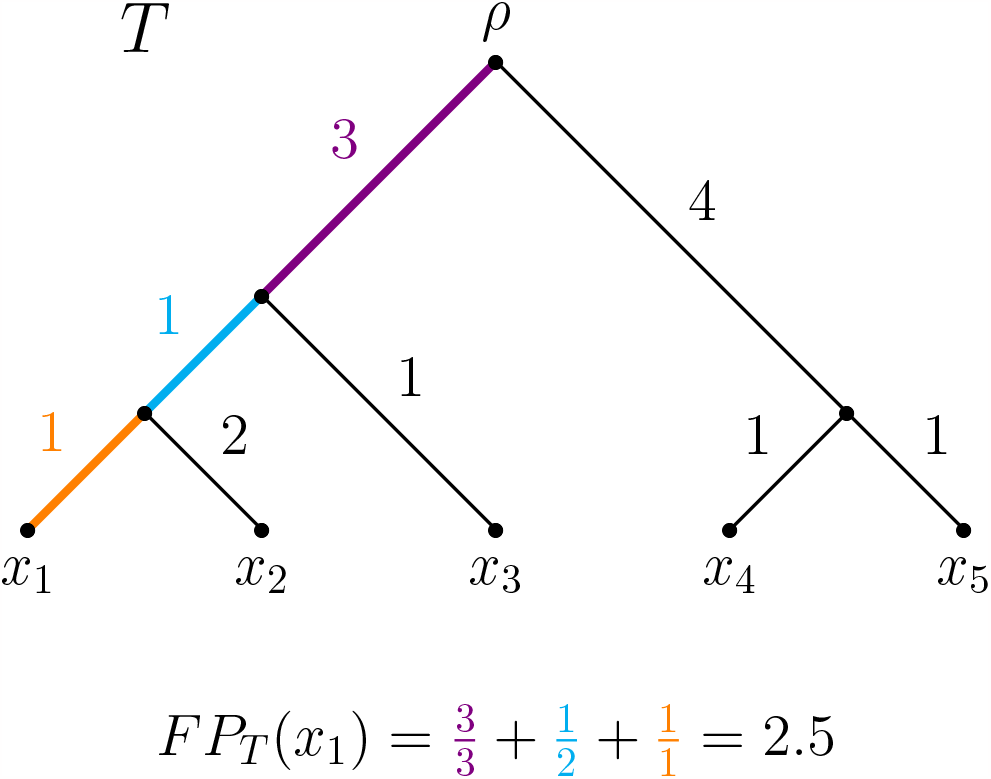
Rooted phylogenetic tree *T* with leaf set *X* = {*x*_1_, …, *x*_5_} and non-negative lengths assigned to its edges. As an example, for the FP index of *x*_1_, we have *FP*_*T*_ (*x*_1_) = ^3^ + ^1^ + ^1^ = 2.5.

### Data collection

Nine multilocus data sets were curated from the literature. In some cases, we reduced the published data sets to a subset of species and/or genes due to large amounts of missing data or missing outgroups. Detailed information on both the data reduction process as well as the species and loci included in our analyses is available from the Appendix. The full data set was used when not otherwise noted below:

- Dolphin data set [12]: DNA sequence data from 24 genes for 47 aquatic mammals. We reduced the data set to 28 species and 22 genes.
- Fungi data set [25]: Amino acid sequence data, gene tree, and species tree estimates based on 683 genes for 25 individuals of bipolar budding yeasts and four outgroups.
- Mammal data set [26, 27]: DNA sequence data and gene tree estimates based on 447 genes for 33 species of mammals and four outgroups.
- Plant data set [25]: DNA sequence data, gene tree, and species tree estimates based on 363 genes for 48 Lamiaceae and four outgroup species. We reduced the data set to 318 genes for the 52 species.
- Primate data set [1]: DNA sequence data from 52 genes for four primate species.
- Rattlesnake data set [9]: DNA sequence data from 19 genes for 24 individuals of six subspecies of *Sistrurus* rattlesnakes and two outgroup species. We picked one sequence per subspecies and one outgroup sequence (i.e., seven sequences in total), and used 16 of the 19 loci.
- Rodent data set [25]: DNA sequence data, gene tree, and species tree estimates based on 794 genes for 37 rodent species. We reduced the data set to 761 genes for the 37 species.
- Snake data set [20]: DNA sequence data, gene tree, and species tree estimated based on 333 genes for 31 caenophidians and two outgroup species.
- Yeast data set [24]: DNA sequence data from 106 genes for eight yeast species.

### Tree estimation and data analysis

Five of the nine data sets (fungi, mammal, plant, rodent, and snake) contained gene tree estimates, which we subsequently used in the analysis. For the remaining four data sets (dolphin, primate, rattlesnake, and yeast), we estimated gene trees under the GTR+Gamma model using RAxML version 8.2.12 [28].

Except for the fungi and snake data sets, a species tree estimate was obtained using SVDquartets [2] as implemented in the PAUP* package [30] (note that we re-estimated species trees for the rodent and plant data sets instead of using the published species trees since we reduced the corresponding data sets in the gene tree analysis). Finally, maximum likelihood branch lengths were computed for all species trees under the GTR+Gamma model (for all but the fungi data set, for which we used the LG model for amino acid sequence evolution) with the molecular clock enforced using PAUP*.

FP indices on individual gene and species trees as well as average FP indices across gene trees were calculated and the resulting values were transformed into rankings (using the “1224” standard competition ranking for ties). Based on this, boxplots of the distribution of ranks across gene trees were generated for each species (an example is depicted in Fig. 2) and their interquartile ranges (IQRs) were computed. The mean, minimum, and maximum IQR across taxa, standardized by dividing by the number of taxa, was computed. Larger values indicate more variability in the distribution of ranks, and thus ranks that differ more across loci. Kendall’s *τ* rank correlation was calculated between the rankings obtained from pairs of gene trees, and from individual gene trees and the species tree (Fig. 3), as well as from the average FP index and the species tree (Table 1). Finally, for each data set, the set of species ranked in the top quartile on the species tree was computed and compared with the percentage of gene trees supporting the placement of these species in the top quartile (Table 2). All statistical analyses were performed using the R Statistical Software [21].

**Table 1:** Summaries of the species rankings for varying phylogenies. Column 2: Kendall’s *τ* rank correlation between the ranking of species induced by the average FP index across gene trees and that induced by the FP index on the species tree. Columns 3-5: Mean, minimum, and maximum IQR of the ranks obtained in the gene tree analyses (standardized by dividing by the number of taxa).

| Data set | Kendall’s $\tau$ | mean IQR | min IQR | max IQR |
| --- | --- | --- | --- | --- |
| Dolphin | 0.8465 | 0.2851 | 0.0000 | 0.5804 |
| Fungi | 0.6651 | 0.2206 | 0.0690 | 0.5690 |
| Mammal | 0.6041 | 0.1651 | 0.0270 | 0.2703 |
| Plant | 0.4828 | 0.1713 | 0.0192 | 0.2308 |
| Primate | 0.5477 | 0.3750 | 0.2500 | 0.5000 |
| Rattlesnake | 0.1502 | 0.3214 | 0.1786 | 0.4643 |
| Rodent | 0.5920 | 0.3084 | 0.0270 | 0.5135 |
| Snake | 0.6298 | 0.2360 | 0.0000 | 0.3939 |
| Yeast | 0.7638 | 0.1875 | 0.1250 | 0.5000 |

**Table 2:** Set of species ranked among the top 25 % of species together with the percentage of gene trees supporting a ranking position in the top quartile for each data set. Note that when the top 25% of species was not an integer (e.g., when the total number of species was not a multiple of four or when there were ties), we extended the set to include some additional species.

| Data set | Top quartile species<br>(species tree) | Percentage of gene trees supporting<br>placement in top quartile (rounded) |
| --- | --- | --- |
| Dolphin | *P. macrocephalus* | 100 % |
|  | *N. phocaenoides* | 68 % |
|  | *P. phocoena* | 86 % |
|  | *D. leucas* | 68 % |
|  | *M. monoceros* | 59 % |
|  | *L. acutus* | 59 % |
|  | *O. orca* | 50 % |
|  | *L. albirostris* | 59 % |
| Fungi | *C. jadinii* | 24 % |
|  | *W. anomalus* | 72 % |
|  | *K. marxianus* | 95 % |
|  | *S. cerevisiae* | 98 % |
|  | *H. gamundiae* | 87 % |
|  | *H. singularis* | 86 % |
|  | *H. jakobsenii* | 93 % |
|  | *H. hatyaiensis* | 60 % |
| Mammal | *G. gallus* | 79 % |
|  | *O. anatinus* | 94 % |
|  | *M. eugenii* | 52 % |
|  | *M. domestica* | 64 % |
|  | *S. tridecemlineatus* | 8 % |
|  | *C. porcellus* | 61 % |
|  | *S. araneus* | 85 % |
|  | *E. europaeus* | 76 % |
|  | *T. belangeri* | 21 % |
|  | *E. telfairi* | 85 % |
| Plant | *A. pectinata* | 99 % |
|  | *L. tibetica* | 10 % |
|  | *P. volubilis* | 90 % |
|  | *S. baicalensis* | 79 % |
|  | *R. myricoides* | 88 % |
|  | *L. angustifolia* | 86 % |
|  | *P. bambusetorum* | 0 % |
|  | *H. sanguinea* | 0 % |
|  | *T. canadense* | 81 % |
|  | *P. lasianthos* | 44 % |
|  | *P. cablin* | 87 % |
|  | *A. reptans* | 71 % |
|  | *C. bungei* | 69 % |
| Primate | *P. pygmaeus* | 23 % |
| Rattlesnake | *A. contortrix* | 81 % |
|  | *S. c. catenatus* | 38 % |

Table 2: *Continued from previous page*
|  |  |  |
| --- | --- | --- |
| Rodent | *R. norvegicus* | 1 % |
|  | *R. fuscipes* | 13 % |
|  | *M. musculus* | 99 % |
|  | *C. gliroides* | 91 % |
|  | *H. minahassae* | 84 % |
|  | *A. imitator* | 38 % |
|  | *L. nouhuysi* | 84 % |
|  | *C. ruemmleri* | 48 % |
|  | *M. major* | 37 % |
|  | *P. macrourus* | 64 % |
|  | *H. goliath* | 48 % |
| Snake | *A. carolinensis* | 98 % |
|  | *P. molurus* | 38 % |
|  | *A. granulatus* | 93 % |
|  | *X. javanicus* | 95 % |
|  | *P. carinatus* | 84 % |
|  | *G. prevostiana* | 46 % |
|  | *L. inornatus* | 14 % |
|  | *P. notostictus* | 51 % |
|  | *P. cana* | 10 % |
|  | *P. frontalis* | 24 % |
| Yeast | *C. albicans* | 93 % |
|  | *S. kluyveri* | 47 % |

**Figure 2:**
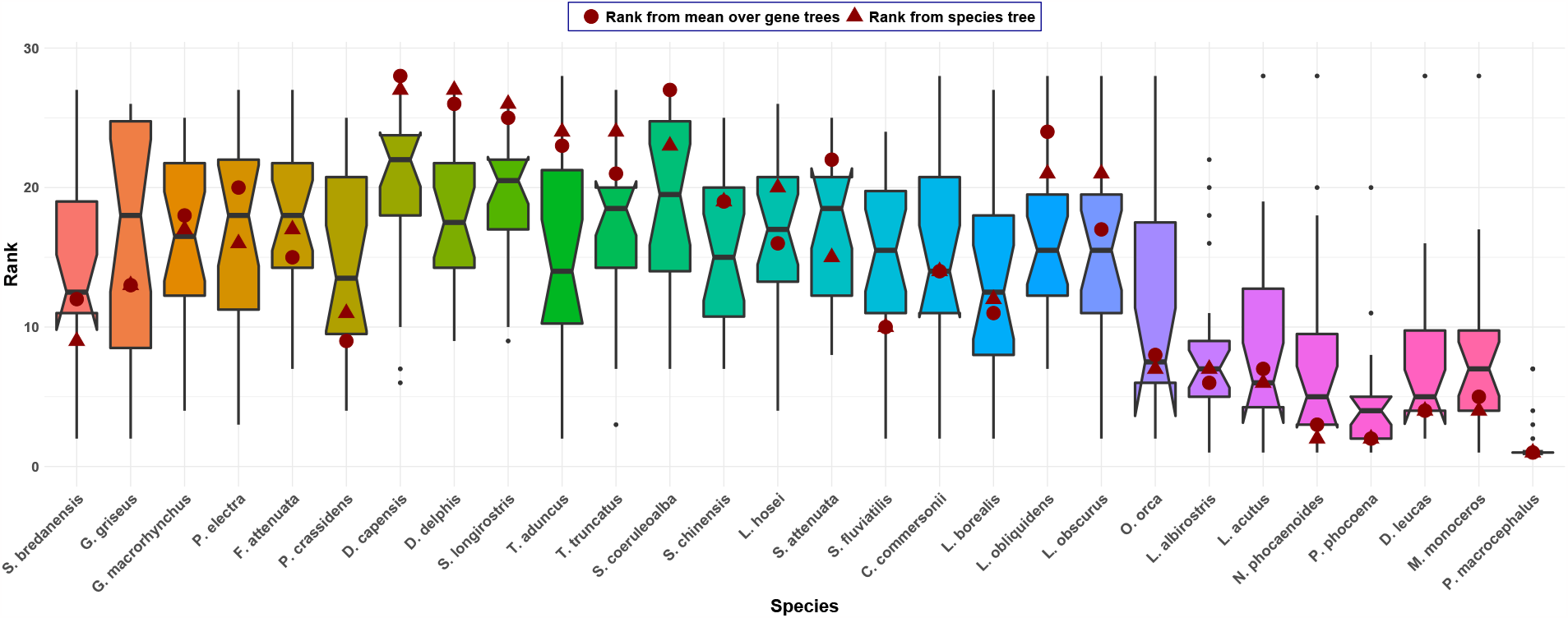
Boxplot of the ranks obtained from the FP index for the Dolphin data set consisting of 28 species and 22 genes. In addition, the ranks obtained from the average FP index across the 22 gene trees (dots) and the ranks on the species tree (triangles) are depicted. The boxplots show large variability in ranks across gene trees. Moreover, the ranks obtained from the average FP index across gene trees and the FP index on the species tree are sometimes ordered differently. For example, *F. attenuata* receives a higher rank than *P. electra* on the species tree, but a smaller rank based on the average FP index.

**Figure 3:**
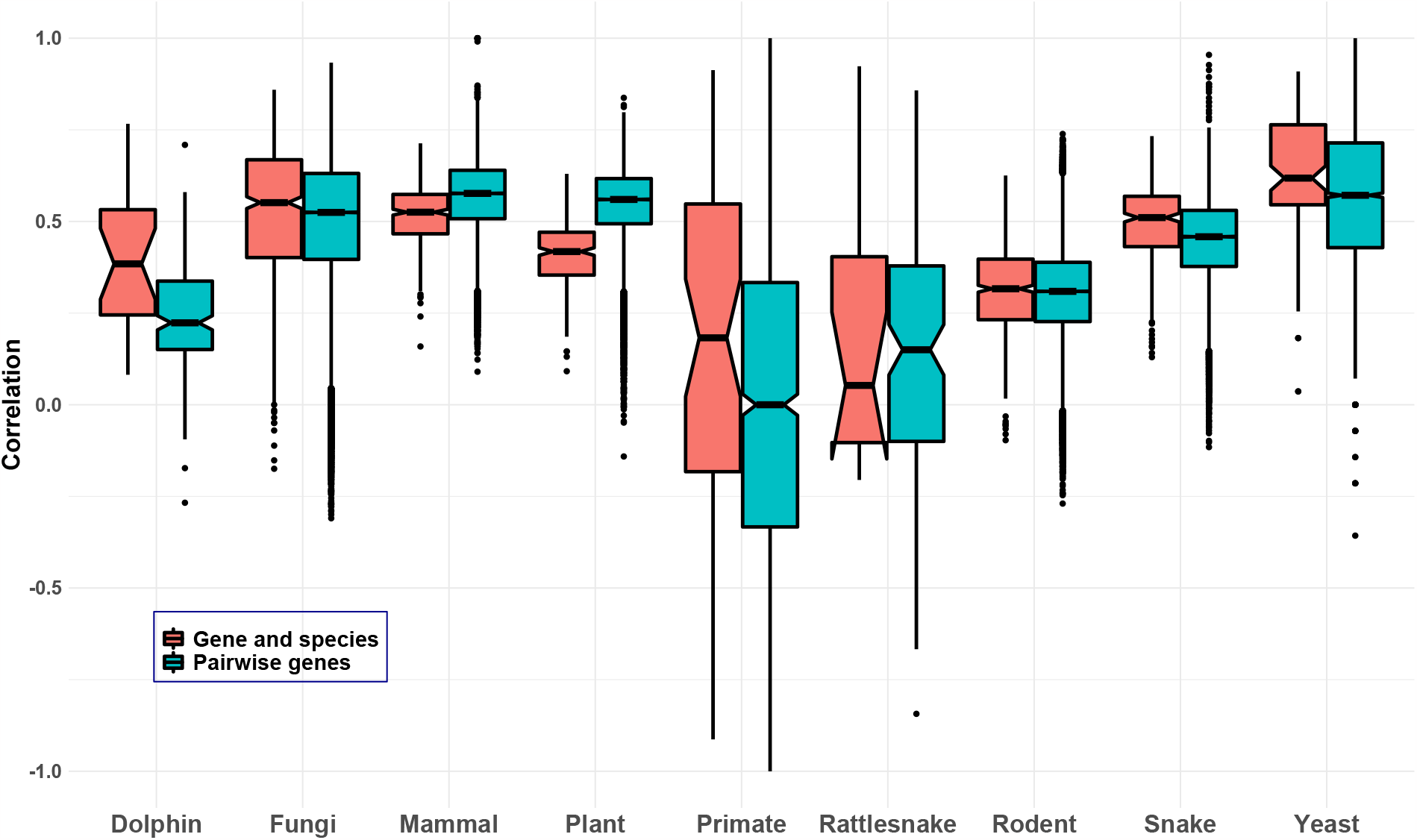
Boxplots of Kendall’s *τ* rank correlation between rankings obtained from individual gene trees and the species tree (red) and rankings obtained from pairs of gene trees (blue) for all data sets considered. Note that for the plant data set, Kendall’s *τ* rank correlation between the rankings obtained from pairs of gene trees was based on the ranks of all species present in both gene trees, respectively (due to missing data, not all gene trees contained the complete set of species).

## Results

We observed extensive variation in the rankings of taxa induced by the FP index across different gene trees for all data sets analyzed. As a representative example, Fig. 2 shows boxplots of the rankings for each of 28 marine mammal species for a data set with 22 genes [12]. Note in particular that some species (e.g., *G. griseus*) have ranks that span almost the entire range of possibilities, while relatively few have consistent ranks across all genes (as would be indicated by “narrow” boxplots in Fig. 2). The corresponding boxplots for the other data sets, which vary in the type of organism and the number of genes included, show similar behavior (see the Supplemental Figures).

To measure the extent of variation in the rankings of species across genes trees, the IQR of ranks assigned was computed for each species. The mean, minimum, and maximum of the IQRs across species (Table 1, columns 3-5) are relatively stable, with the mammal, plant, and yeast data sets showing the lowest variability in rankings and the primate data set the highest. The large maximum IQR for the dolphin and fungi data sets suggests the presence of at least one species for which large variation in ranks across genes is observed.

Fig. 3 summarizes results across all data sets examined. The blue boxplots show the distribution of values of Kendall’s *τ* rank correlation between rankings of species obtained from pairs of gene trees. While there tends to be a positive correlation overall, some boxes are closer to zero than to one, and some negative values (meaning that the corresponding rankings are negatively correlated; in other words, they tend to rank species in opposite order) are present. Thus, using different gene trees for the calculation of the FP index can result in very different prioritization orders.

A similar trend occurs for the correlation between rankings obtained from individual gene trees and the species tree. The red boxplots in Fig. 3 again show a tendency for positive correlations, but in several cases the correlation is closer to zero than to one, and for some data sets (e.g., the primate and rattlesnake data sets) negative values are again observed.

Given the variability in ranks obtained from different gene trees, we additionally considered how the FP index averaged across gene trees compared to that obtained from the species tree. For most data sets, we observe some changes in the orderings of ranks when using the average over gene trees vs. the species tree (see Fig. 2 for an example). However, as Table 1 shows, the rankings are all positively correlated and, with the exception of the plant and rattlesnake data sets, the correlation is larger than 0.5. Thus, the average FP index across loci and the FP index based on the species tree tend to result in similar prioritization orders of species in general.

Finally, from a conservation view point it may be relevant to know whether e.g., the top 25% of species are consistently placed in the top quartile of species across different gene trees, even if there are small rank switches among them. For this reason, we additionally computed the set of species placed in the top quartile in the ranking based on the species tree, together with the percentage of gene trees supporting its placement in the top quartile. Table 2 shows that in most cases species ranked high on the species tree are indeed also ranked high on the majority of gene trees. This indicates that while there might be small differences in the precise ranking orders, different gene trees mostly agree on the set of highly ranked species. However, there are some outliers with very low support on the gene tree level. These correspond to species for which the rank on the species tree is also very different from the average rank across gene trees (see, e.g., *P. bambusetorum* and *H. sanguinea* in the plant data set and compare this to the rank from the species tree and the rank from the mean over gene trees given in the Supplemental Figures).

## Discussion

Many questions in evolutionary biology require the specification of a phylogeny as a precursor to downstream analyses and are thus challenged by conflicting evolutionary signal such as hetereogeneity in gene trees or discordance between gene trees and the species tree. In this case study, we employed the FP index, a popular tool for prioritizing species for conservation based on their placement in an underlying phylogeny, to illustrate the effect of gene tree variation on downstream phylogenetic analyses. In analyzing species rankings obtained from the FP index for nine multilocus data sets for a broad range of organisms, we found that different loci/gene trees can induce very different prioritization orders of species (reflected by large variability in the rank a species is assigned across gene trees). However, with few exceptions, taking all loci into account by averaging the FP index across loci or by constructing a species phylogeny based on all loci resulted in similar prioritization orders. Additionally, we found that the set of species ranked in the top quartile on the species tree is in most cases also ranked highly by the majority of gene trees. On one hand, this result is in line with the previous observation that averaging the FP index across loci tends to result in similar prioritization orders as using the species tree. On the other hand, it may be relevant for actual conservation action targeted at the top set of species rather than individual species, as it illustrates that while different gene trees may rank species differently, in most cases they tend to agree on the set of most evolutionary distinct species. However, we also saw some exceptions to this trend, i.e., species ranked highly on the species tree, but with very low support on the gene tree level, and it would be interesting to investigate the reasons and implications of this further in future research.

While we have used the FP index to illustrate the effect of gene tree variation on prioritization, this phenomenon is not limited to the FP index. Because phylogenetic diversity indices rely on specification of an underlying phylogeny, all such indices have the potential for the particular phylogeny chosen to strongly influence the results. We remark, however, that the FP index is weighted towards the tips of the phylogeny (since, by its definition, the lengths of edges close to the tips have a higher influence than more ancient edges) and thus may be more strongly affected by processes like incomplete lineage sorting than other PD indices. It would thus be interesting to see how other PD indices, such as the new ED2 score mentioned in the introduction, fare in this setting.

More generally, it would be interesting to analyze the behavior of other phylogenetic downstream analyses in the presence of phylogenetic heterogeneity. Given the role of evolutionary processes like incomplete lineage sorting and hybridization in generating variation in the trees for individual loci throughout the genome, the choice of phylogeny to use for downstream analyses is far from straightforward. Recent work has for instance shown that analysis of quantitative trait evolution can benefit from the incorporation of gene tree variation [13]. The analyses presented here thus serve to underscore the importance of carefully considering the choice of phylogeny for settings such as these.

## Conclusion

Our results demonstrate a need for future research into the effect of phylogeny choice on downstream inference. While in many cases use of a species tree might provide a robust estimate of evolutionary relationships, an argument can be made that use of a species tree could obscure relationships of interest to a particular question if such relationships were strongly supported only by a subset of genes. Approaches that more explicitly accommodate variability in the underlying histories of the loci by computing single-locus trees may be promising in such cases, as they would allow an examination of the contribution of individual phylogenies to the overall results. However, this comes at the expense of increased computation, as such approaches would require separate estimation of phylogenies for all individual loci. Another possibility would be to adopt a Bayesian approach to formally integrate over the distribution of gene trees, again incurring a substantial computational cost. Overall, we hope that our analyses of the FP index provide a call for future research on how to best infer evolutionary relationships for downstream analyses.

## Supplementary information

Additional information and figures can be found in the Supplementary Information (SI) available at https://github.com/lkubatko/FP-Index-Discordance.

## Acknowledgments

We thank two anonymous reviewers of an earlier version of this paper for their valuable feedback and suggestions.

